# Septins mediate a microtubule-actin crosstalk that enables actin growth on microtubules

**DOI:** 10.1101/2022.02.15.480618

**Authors:** Konstantinos Nakos, Megan R. Radler, Ilona A. Kesisova, Meagan R. Tomasso, Shae B. Padrick, Elias T. Spiliotis

**Affiliations:** Department of Biology, Drexel University, Philadelphia, PA 19104; Department of Biochemistry and Molecular Biology, Drexel University, Philadelphia, PA 19102

**Keywords:** septins, actin filaments, microtubules, actin microtubule crosstalk, growth cones, neurons

## Abstract

Cellular morphogenesis and processes such as cell division and migration require the coordination of the microtubule and actin cytoskeletons (1, 2). Microtubule-actin crosstalk is poorly understood and largely regarded as the capture and regulation of microtubules by actin (1, 2). Septins are filamentous GTP-binding proteins, which comprise the fourth component of the cytoskeleton along microtubules, actin and intermediate filaments (3, 4). Here, we report that septins mediate microtubule-actin crosstalk by coupling actin polymerization to microtubule lattices. Super-resolution imaging shows that septins localize to overlapping microtubules and actin filaments in the growth cones of neurons and non-neuronal cells. We show that recombinant septin complexes directly crosslink microtubules and actin filaments into hybrid bundles. In vitro reconstitution assays reveal that microtubule-bound septins capture and align stable actin filaments with microtubules. Strikingly, septins enable the capture and polymerization of growing actin filaments on microtubule lattices. In neuronal growth cones, septins are required for the maintenance of the peripheral actin network that fans out from microtubules. These findings provide the first evidence of septins directly mediating microtubule interactions with actin filaments, and reveal a new mechanism of microtubule-templated actin growth with broader significance for the self-organization of the cytoskeleton and cellular morphogenesis.

**Significance Statement:** Cellular morphogenesis and processes such as cell division and cell migration require the coordination of the actin and microtubule cytoskeletons. Despite its broader physiological significance, actin-microtubule crosstalk is molecularly and mechanistically poorly understood. Actin-microtubule crosstalk has been viewed primarily as the regulation or guidance of microtubule dynamics by actin filaments. Here, we have discovered a new mechanism by which actin filament growth is guided by microtubules. We report that septins, a poorly understood component of the cytoskeleton, can capture and link the polymerizing ends of actin filaments to microtubule polymers. We present evidence that this new septin-mediated mechanism is critical for the morphology of growth cones, which are structures that direct the growth of neuronal axons through actin-microtubule crosstalk.

## Introduction

Cell morphology, dynamics and processes such as mitosis and migration require crosstalk between the actin and microtubule cytoskeletons (1, 2). Actin-microtubule crosstalk encompasses the physical crosslinking of actin and microtubule polymers, and the capture, hindrance or guidance of polymerizing microtubules by or on actin filaments (1, 2). Regulation of actin organization and dynamics by microtubules is a type of crosstalk, which is less evident and studied. Microtubules regulate actin organization indirectly by harboring signaling factors, but recent studies indicate that actin can polymerize directly from microtubule plus ends (1, 5, 6). Microtubule-based nucleation of actin filaments might be critical for the steering of the growth cones of neuronal axons, which depends on microtubules, and microtubule-based protrusions (7, 8). Overall, however, microtubule-dependent actin growth remains mechanistically and physiologically poorly understood.

Septins are a family of GTP-binding proteins that assemble into higher order oligomers and polymers, which comprise the fourth component of the cytoskeleton after actin, microtubules and intermediate filaments (3). Septins associate with subsets of actin filaments and microtubules (4). Septins have been shown to directly crosslink actin filaments or microtubules into bundles, but it is unknown if septins physically interact with both actin and microtubules, linking their dynamics and organization (4). Here, we report that septins provide a novel mechanism of actin-microtubule crosstalk, mediating the capture and growth of actin filaments on microtubule lattices, and show that septins are an essential component of the actin-microtubule coordination that maintains the morphodynamics of neuronal growth cones.

## Results and discussion

Neuronal growth cones are an ideal model system for studies of actin-microtubule crosstalk (1, 2, 7). At the tip of axons and dendrites, growth cones have a fan-like morphology that consists of a central microtubule-containing domain, which is proximal to the axon shaft, and a peripheral actin-rich domain with lamellipodial and filopodial protrusions (7). Growth cone morphology and dynamics depend largely on the crosstalk between the peripheral actin network and microtubules that invade from the central into the peripheral domain (7). We, therefore, sought to examine the localization of septins in the growth cones of primary rat hippocampal and differentiated B35 neurons using high- and super-resolution microscopy. We stained neurons for Sept7 as a common subunit and marker of septin complexes. In the peripheral domain of growth cones, fluorescence line scans showed that Sept7 colocalizes with microtubule segments that coalign with actin bundles (Fig. 1A-B). Super-resolution structured illumination showed that septin filaments adjoin actin and microtubules at the base of filopodia, and septin puncta localize on points of contact between the distal ends of microtubules and the proximal ends of filopodial actin bundles (Fig. 1C). Sept7 also coaligns with actin and microtubules in filopodia (Fig. 1D), and localizes at the orthogonal junctions of axonal shaft microtubules with filopodial actin filaments (Fig. 1E). Super-resolution imaging of the kidney fibroblast COS-7 cells showed that coalignment of microtubules with actin filaments and septins is conserved across different cell types (Fig. 1F).

**Fig. 1.**
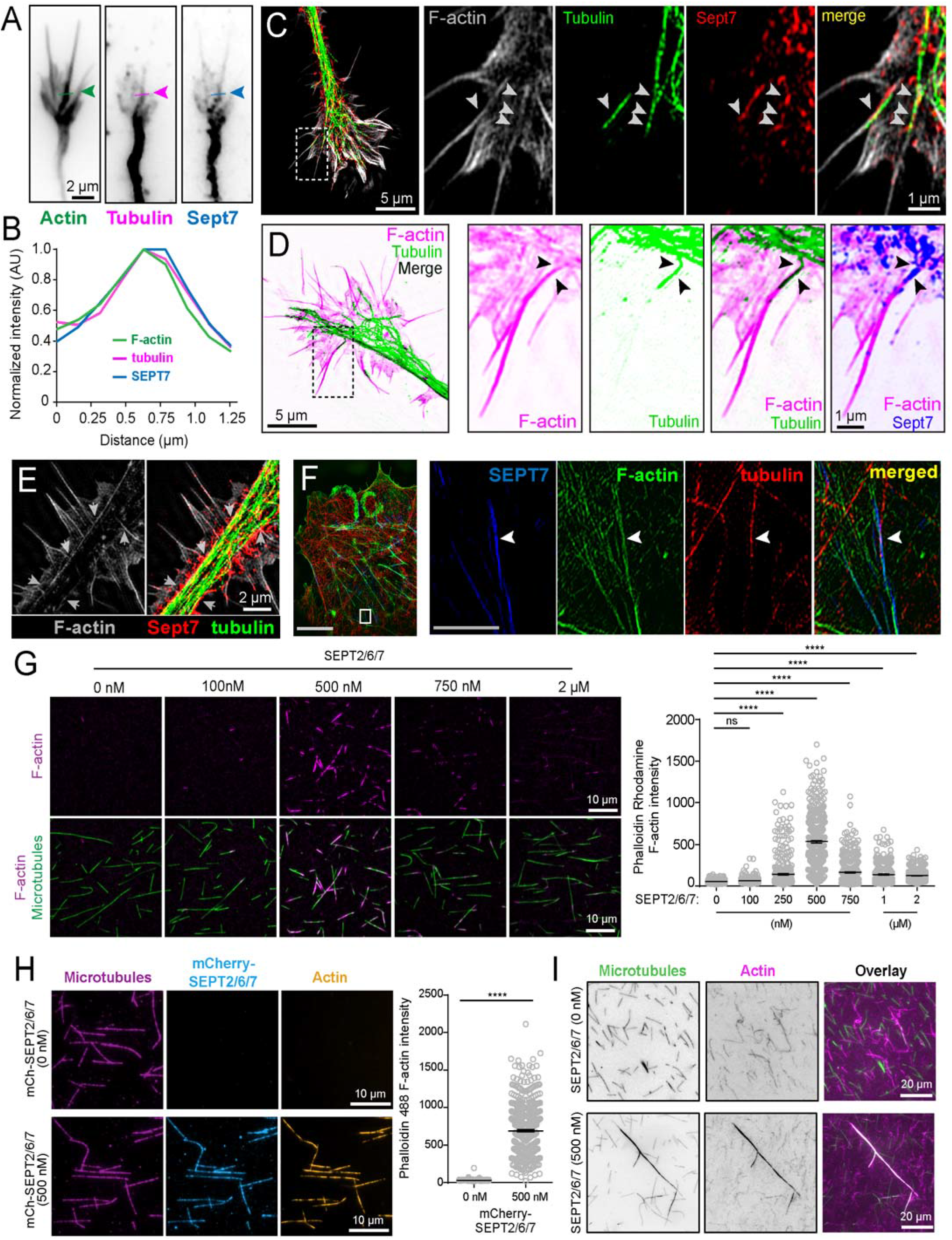
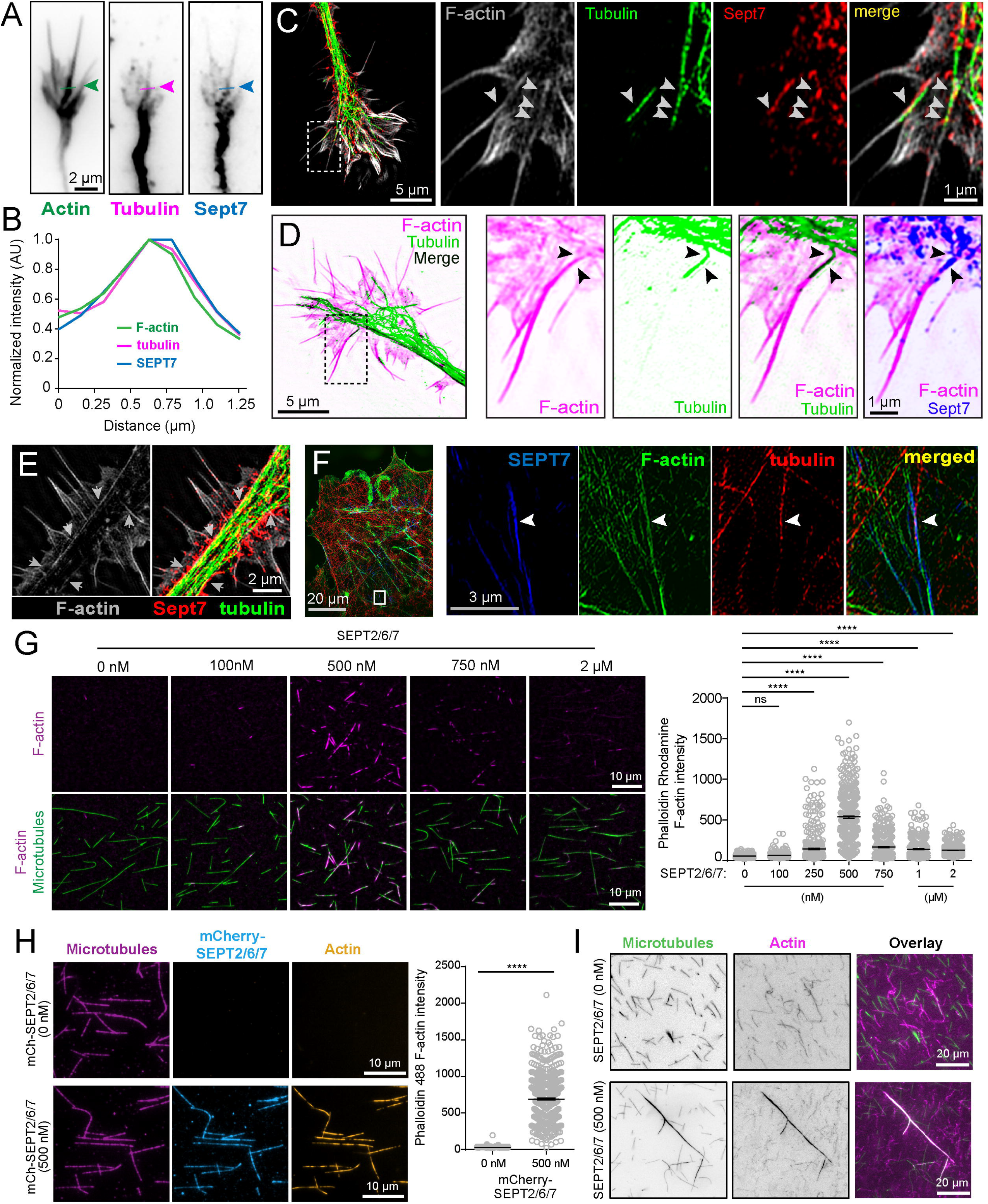
Septins colocalize with overlapping microtubules and actin filaments, and mediate actin-microtubule binding. (A-B) Images show the axonal growth cone of a rat hippocampal neurons (DIV6) stained for F-actin (phalloidin), α-tubulin and Sept7. Line scans shows fluorescence overlap between a microtubule, F-actin and Sept7 in the peripheral domain of the growth cone. (C-E) Super-resolution SIM images of the axonal growth cone of differentiated B35 neurons, which were stained for F-actin (phalloidin), α-tubulin and Sept7. Arrowheads (C-D) point to overlapping F-actin, microtubules and Sept7, and arrows (E) point to Sept7 at the orthogonal junctions of filopodial F-actin bundles with axon shaft microtubules and a coaligning actin cable. (F) SIM image of a COS-7 cell shows coalignment (arrowhead) of a microtubule (red) with Sept7 (blue) and actin filaments (red). (G) TIRF images of phalloidin-stabilized actin filaments (magenta), which were flowed into a chamber with immobilized microtubules (green) after incubation with increasing Sept2/6/7 concentrations. Plot shows the average intensity of microtubule-bound actin after 15 minutes of incubation with taxol-stabilized microtubules, which were decorated with 0 nM (*n* =424), 100 nM (*n* =334), 200 nM (*n* =406), 500 nM (*n* =455), 750 nM (*n* =433), 1 μM (*n* =391) and 2 μM (*n* =371) of Sept2/6/7. (H) TIRF images of taxol-stabilized microtubules, which were decorated with mCherry-Sept2/6/7 (0 or 500 nM) and subsequently incubated with phalloidin-stabilized actin filaments for 15 minutes. Plot shows the average intensity of microtubule-bound actin in 0 nM (*n* =656) and 500 nM (*n* =636) mCherry-Sept2/6/7. (I) Images show the result from mixing in solution pre-polymerized phalloidin-stabilized actin filaments with taxol-stabilized microtubules in the absence or presence of Sept2/6/7. n.s., non significant (p > 0.05); ***, p < 0.0001.

To test whether septins interact directly and concomitantly with both actin and microtubules, crosslinking them together, we performed in vitro binding assays using stable prepolymerized microtubules, actin filaments and septins. We purified recombinant Sept2/6/7, the minimal septin complex which has been shown to bind separately actin and microtubules (9, 10). Using total internal reflection fluorescence (TIRF) microscopy, we assayed for the association of immobilized glass-bound Sept2/6/7-coated microtubules with actin filaments that were floated into the imaging chamber. In the physiological range of intracellular septin concentrations (200-800 nM) (11), actin filaments bound microtubules in a septin concentrationdependent manner (Fig. 1G). Actin-microtubule binding peaked at 500 nM of Sept2/6/7, and tapered off with higher concentrations (Fig. 1G). This biphasic effect resembled the increase and diminution of microtubule plus end growth by nanomolar and micromolar Sept2/6/7 concentrations, respectively, which corelates with a transition of Sept2/6/7 from oligomers to higher-order polymers with increasing concentrations (9). Coating microtubules with mCherry-Sept2/6/7 showed that septins are present along the actin-bound microtubule lattice (Fig. 1H). Pre-binding of Sept2/6/7 to microtubules was not required for actin-microtubule crosslinking. Mixing Sept2/6/7 with pre-polymerized stable microtubules and actin filaments in solution resulted in elongated bundles consisting of both microtubules and actin (Fig. 1I). Taken together with the intracellular localization of septins to actin-bound microtubules, these data demonstrate that septins mediate actin-microtubule interactions and therefore, can facilitate actin-microtubule crosstalk.

We next sought to examine if septins mediate interactions between polymerizing actin filaments and microtubules. Previously, we found that Sept2/6/7 filaments can transiently pause and capture polymerizing microtubule plus ends, but it is unknown whether septins affect similarly the fast-growing ends of actin. We reconstituted actin growth from immobilized phalloidin-stabilized actin seeds in vitro, and imaged their interaction with microtubules in the presence or absence of Sept2/6/7. We found that Sept2/6/7 increased the overlap between microtubules and polymerizing actin filaments by two-fold (Fig. 2A-B). Analysis of the collision events between growing actin ends and microtubules showed that in the absence of Sept2/6/7, actin ends crossed over microtubules with virtually no overlapping or zippering (Fig. 2C; Movie S1). On Sept2/6/7-coated microtubules, however, there was a drastic increase in the overlap of polymerizing actin ends with microtubules at collisions of <30° angle (Fig. 2C). Strikingly, polymerizing actin ends grew along the lattice of microtubules in a zippering manner, and continued to grow after overtaking the distal microtubule end with the preceding polymerized filament remaining bound to the microtubule (Fig. 2D; Movies S2 and S3). Notably, the rate of actin elongation did not change upon attachment to and zippering along the Sept2/6/7-coated microtubule lattice (1 ± 0.03 vs 0.97 ± 0.04 μm/minute; *n* = 13), which suggests that microtubule-bound Sept2/6/7 associates with polymerizing actin filaments through weak and/or transient interactions that do not sterically hinder actin growth. Thus, septins can mediate a microtubule-templated actin growth, capturing polymerizing actin ends and facilitating their growth along microtubule lattices.

**Fig. 2.**
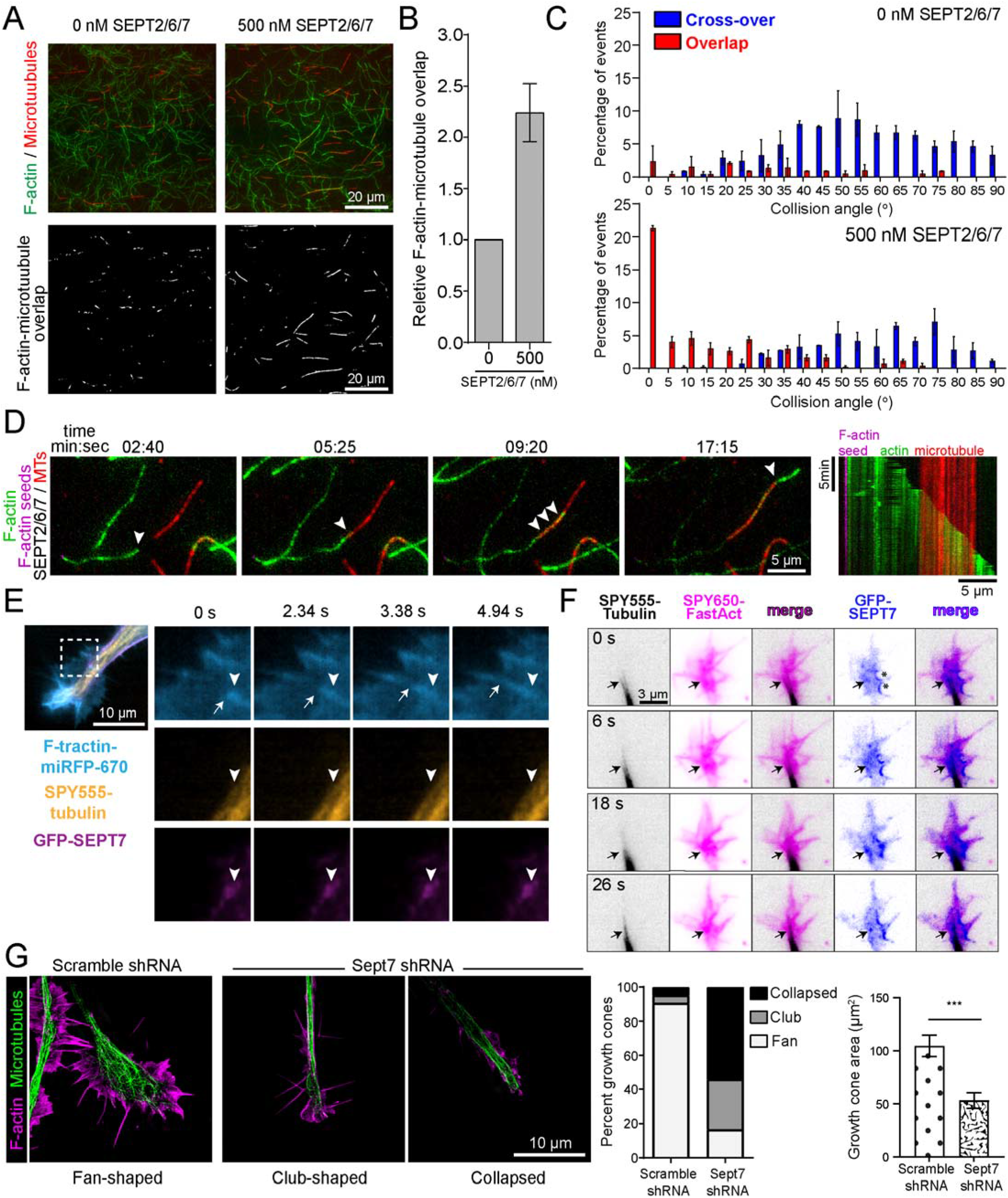
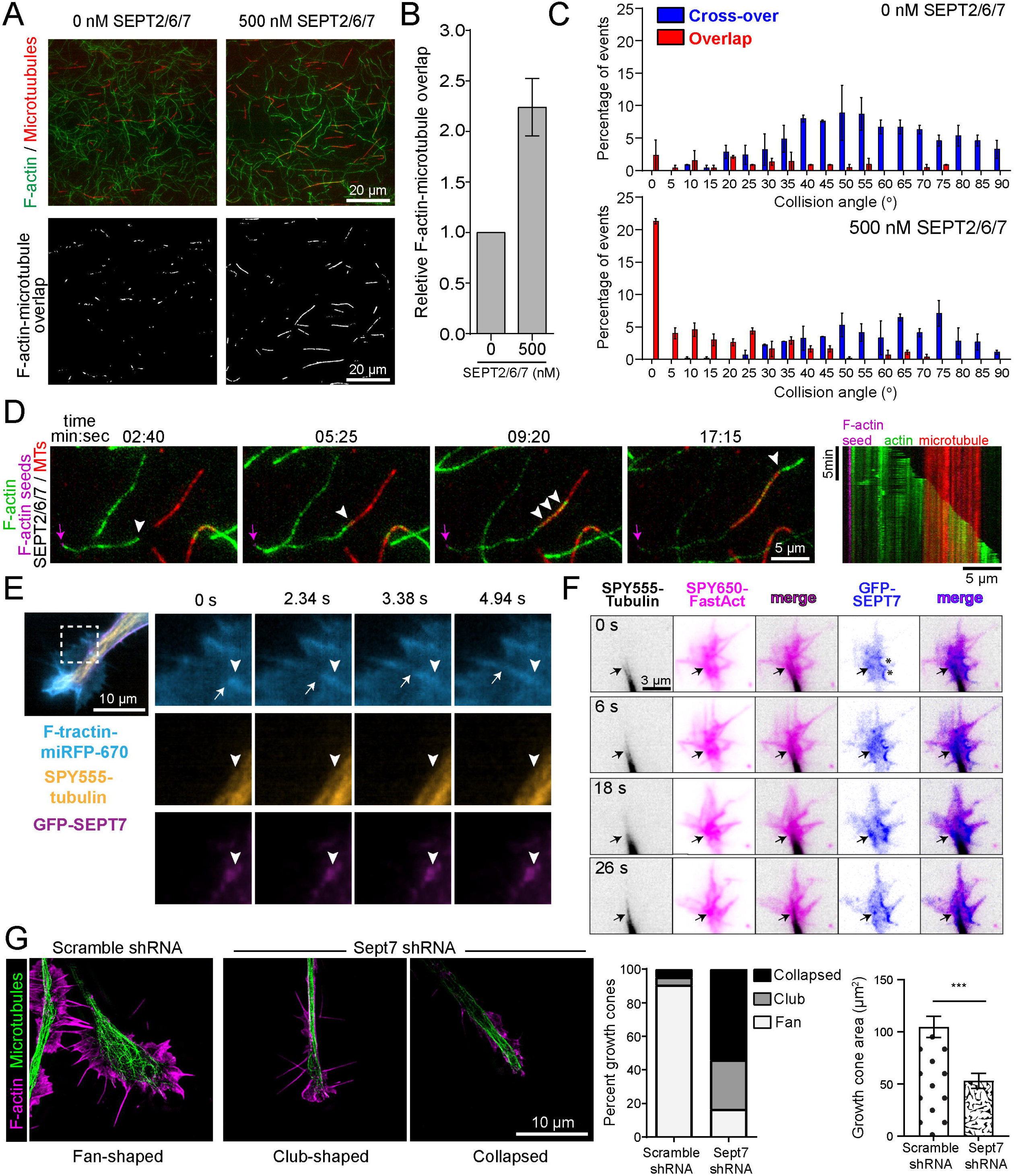
Septins enable actin polymerization on microtubule lattices and required for protrusive growth cone morphology. (A) TIRF images show actin filaments (green) and microtubules (red), and their overlap (grey) after 30 minutes of actin polymerization from immobilized actin seeds in the presence of taxol-stabilized microtubules, which were coated with Sept2/6/7. (B) Bar graph shows the relative increase in the microtubule-bound actin as a fraction of total actin, which was measured as the overlapping over the total surface area of actin filaments. (C) Plots show the distribution of the angles of collision between polymerizing actin ends and microtubules as percentage of total collisions, and their categorization into overlapping (red) or non-overlapping crossover (blue) events based on actin end coalignment with the microtubule lattice or lack thereof (*n* = 236-397 events). (D) Still frames and kymograph of time-lapse TIRF imaging of the capture and zippering of a polymerizing actin end (arrowhead) along a stable Sept2/6/7-coated microtubules. Arrow points to the actin seed. (E) Still frames from time-lapse TIRF imaging of growth cones of living B35 neurons, which expressed GFP-Sept7 and F-tractin-miRFP670, and were labeled with SPY555-tubulin. Arrowhead and arrow point to GFP-Sept7 and a polymerizing actin filament, respectively. (F) Time-lapse TIRF image of the axonal growth cone of a rat hippocampal neuron (DIV3) that was transfected with GFP-Sept7 and labeled with SPY555-tubulin and SPY650-FastAct. Arrows point to GFP-Sept7 overlap with microtubules and actin. Asterisks denote GFP-Sept7 localization to areas of membrane curvature. (G) Images show representative fan- and club-shaped, and collapsed growth cones of differentiated B35 neurons, which were stained for microtubules (α-tubulin; green) and F-actin (phalloidin; magenta) after transfection with control and Sept7 shRNAs. Bar graph shows the percentage distribution of growth cones (*n* = 21-24) based on their morphology, and their mean (± SEM) surface area in neurons. ***, *p* < 0.0001.

Capture and growth of polymerizing microtubules along actin has been the predominant mode of actin-microtubule crosstalk in vivo. Surprisingly, in vitro assays showed that actin can polymerize directly from microtubule plus ends (5), and recent work indicates that the microtubule plus end protein adenomatous polyposis coli (APC) promotes actin polymerization at focal adhesions and neuronal growth cones (6, 12). Our results suggest that actin growth could also occur on microtubule lattices. To probe for events of actin growth on microtubule lattices in living cells, we imaged actin and microtubule dynamics in the growth cones of differentiated B35 and primary rat hippocampal neurons that expressed GFP-Sept7. In the central domain of neuronal growth cones, we observed actin filaments growing from the sides of microtubules in an angled orientation (Fig. 2E, arrows). These growth events took place on microtubules that were fully embedded in the veil of the growth cone, and originated from GFP-Sept7 puncta (Fig. 2E, Movie S4). In addition, GFP-Sept7 localized to actin-rich areas that overlapped with the distal ends of microtubules that protruded from the central into the transitional and peripheral domains of growth cones (Fig. 2F, arrows). In parallel, GFP-Sept7 also decorated saddle-shaped domains of membrane curvature (Fig. 2F, asterisks), which was consistent with septin enrichment on membrane domains of micron-scale curvature (13), indicating that GFP-Sept7 was expressed and localized at physiological levels.

Given that the morphodynamics and protrusive activity of growth cones depend on actin-microtubule crosstalk, we reasoned that septins may impact growth cone morphology. In differentiated B35 neurons, Sept7 depletion shifted the distribution of growth cones from fan-to club-shaped and fully collapsed, which phenotypically resembles the response to repulsive cues that cause growth cone retraction (Fig. 2G). Quantification of growth cone areas and number of filopodia per growth cone showed a ~50% reduction in Sept7-depleted neurons (Fig. 2G). In primary hippocampal neurons, there was similar decrease in fan-shaped growth cones of all neurites from 45% to 20%, and mean surface area was reduced from 12.7 ± 1.4 μm^2^ to 8.2 ± 1.1 μm^2^ (*n* = 174-182 neurites from 20 neurons; *p* = 0.0001). These data reveal a hitherto unknown function for septins in the maintenance of protrusive growth cones.

In sum, our results report the first evidence of septins directly mediating actin-microtubule interactions, enabling actin growth on microtubules, and promoting a protrusive growth cone phenotype. Recent work has unexpectedly revealed that linear and branched actin can grow directly from microtubule plus ends through formins and the adenomatous polyposis coli (APC), which induces actin-driven membrane protrusions in neuronal growth cones (5, 6). Here, the discovery of a novel septin-mediated mechanism of actin polymerization along microtubules indicates that microtubule lattices can guide or template actin polymerization in a similar fashion to how actin filaments provide tracks for microtubule plus end growth. We posit that this mechanism might more widely utilized than hitherto reported; in vitro evidence suggests tau and the growth arrest-specific 2-like (Gas2L1) can couple polymerizing actin to microtubules (14, 15). Capture of polymerizing actin on the surface of microtubules could be of key importance in the trafficking and positioning of membrane organelles (e.g., endosomes, Golgi), which nucleate actin or are bound to actin densities, and their transitioning from microtubule-to actin-dependent transport.

## Materials and Methods

Primary neurons were isolated from freshly dissected embryonic (E18) rat hippocampus (BrainBits/Transnetxy Tissue), and B35 rat neuroblastoma cells were differentiated in Dulbecco’s Modified Eagle (DME) media containing 0.1 mM dbcAMP (Sigma-Aldrich) and N2 supplement (ThermoFisher Scientific), and plated on 1.25 μg/ml laminin and/or 1 mg/ml poly-L-lysine. COS-7 (ATCC: CRL-1651) cells were maintained in high glucose DME (Sigma) with 10% fetal bovine serum (R&D Systems). Structured illumination and TIRF microscopy were performed on the OMX V4 DeltaVision imaging platform (GE Healthcare) with 60X/1.42 NA and 60X/1.49 NA (Olympus) objectives, respectively. Immunofluorescence staining was performed with rabbit anti-SEPT7 (IBL America), mouse anti-tubulin (DM1a, Sigma), and iFluor488-(ATT Bioquest) or rhodamine-phalloidin (Cytoskeleton Inc). In vitro experiments were performed with rabbit skeletal muscle actin and porcine brain tubulin. Details of all the materials and methods are provided in *SI Appendix*.

## ACKNOWLEDGMENTS

This work was supported by National Institutes of Health grant GM 5 R35 GM136337-02 and a Pennsylvania Department of Health CURE grant SAP 4100085747 to E.T.S. All microscopy imaging was performed at the Cell Imaging Center of Drexel University.

