## Supplemental Information for "Septins mediate a microtubule-actin crosstalk that enables actin growth on microtubules"

**This PDF includes:**

Supplementary Text

SI References

**Other supplementary materials for this manuscript include the following:**

Movie S1

Movie S2

Movie S3

Movie S4

**Supplementary Text**

**Cell Culture**

African green monkey kidney COS-7 cells (ATCC: CRL-1651) were maintained in a humidified incubator at 37 ^o^C with 5% CO_2_ in high glucose Dulbecco’s Modified Eagle (DME) medium (Sigma) supplemented with 10% fetal bovine serum (R&D Systems) and 1% penicillin/streptomycin/kanamycin (Sigma and GIBCO). The rat neuroblastoma B35 cells were maintained in DME medium as previously described (1, 2), and differentiated after 24 h by incubation in media containing 0.1 mM dibutyrl cyclic AMP (dBcAMP; Sigma; D0627) and N2 supplement (ThermoFisher Scientific; 17504044). Primary rat hippocampal neurons were isolated from freshly dissected rat embryonic (E18) hippocampus (BrainBits; Transnetyx Tissue). Hippocampi were incubated for 10 minutes at 30-35 ^o^C in Hibernate E medium without calcium (HE-Ca; BrainBits) supplemented with 2 mg/ml papain, and subsequently triturated for one minute continuously in HE-Ca buffer without papain using a salinized long-stem Pasteur glass pipette, which was previously pulled over a Bunsen burner fire to generate a smaller opening. Neurons were pelleted, counted and plated at a density of 60,000 per 12 mm glass coverslip (round German coverslip No. 1; Bellco Glass) or 150,000 per 22 mm glass coverslip or 35 mm glass bottom dish No. 1.5 (Mattek). Glass coverslips and dishes were coated with laminin (1.25 μg/ml; Sigma L2020) and/or 1 mg/ml poly-L-lysine (1 mg/ml; Peptides International OKK-3056).

**Plasmids**

Plasmids for the bacterial expression of His-SEPT2 (pnEA-vH), His-SEPT2-mCherry (pET15b) and SEPT6/7-strep (pnCS), and the purification of dark or mCherry-Sept2/6/7 complexes were described previously (3, 4). The plasmid pEGFP-C1-SEPT7 was a kind of gift from Dr. Smita Yadav (5). The plasmid encoding for F-tractin-miRFP670 was constructed by removing tdTomato from the plasmid pdt-Tomato-N1 F-tractin (a gift from Dr. Tatyana Svitkina, University of Pennsylvania) (6, 7) and replacing it with miRFP670 from the pmiRFP670-N1 plasmid (a gift from Vladislav Verkhusha, Albert Einstein College of Medicine; addgene plasmid # 79987) (8) using the restriction enzymes AgeI and NotI. ShRNA plasmids were constructed by cloning an shRNA that targets the 3'-UTR of the rat *Sept7* gene (GATAAATTGCCATAATATG) and a non-targeting scrambled sequence (control; ATGACTAGATTGATTACAA) into the BglII and HindIII sites of a p-SUPER GFP plasmid. For the Sept7 and scramble shRNAs, the following oligonucleotides were synthesized: 5'-CCCGATAAATTGCCATAATATGTTCAAGAGA CATATTATGGCAATTTATCTTTTTA-3' and 5'-CCCATGACTAGATTGATTACAATT CAAGAGATTGTAATCAATCTAGTCATTTTTTA-3' with 5'-GATC-3' and 3'-TCGA-5' overhangs.

Plasmids that expressed shRNAs and mCherry instead of GFP were constructed by sub-cloning mCherry into the restriction enzyme sites AgeI and BsrG1 of the pSUPER-GFP plasmid.

**Immunofluorescence microscopy**

Rat hippocampal neurons (DIV6; Fig. 1A) were fixed, stained and imaged with an inverted Zeiss AxioObserver microscope with 63X/1.4NA objective, Hamamatsu Orca R2 CCD camera and the Slidebook 6.0 software as previously described (9). For super-resolution microscopy (Fig. 1C-E and 2G), neurons were fixed in PHEM buffer (60 mM Pipes-KOH, pH 6.9, 25 mM Hepes, 10 mM EDTA, and 2 mM MgCl_2_), 3% paraformaldehyde (Electron Microscopy Sciences) and 4% sucrose for 10 minutes. Cells were quenched with 75 mM NH_4_Cl and simultaneously blocked and permeabilized in GDB/Triton buffer (30 mM sodium phosphate pH 7.4, 0.2% gelatin, 450 mM NaCI, 0.1% Triton X-100) for 55 minutes at room temperature. Subsequently, primary antibodies were diluted in GDB/Triton buffer and incubated overnight at 4°C. Secondary antibodies and phalloidin were diluted in GDB/Triton buffer and incubated at room temperature for 1 hour. COS-7 cells (Fig. 1F) were fixed in PHEM buffer (60 mM Pipes-KOH, pH 6.9, 25 mM HEPES, 10 mM EDTA, and 2 mM MgCl_2_), 3% paraformaldehyde (Electron Microscopy Sciences) for 15 minutes. Cells were quenched with 75 mM NH_4_Cl and permeabilized in GDB/Triton buffer (30 mM sodium phosphate pH 7.4, 0.2% gelatin, 450 mM NaCI, 0.1% Triton X-100) for 10 minutes at room temperature. Subsequently cells were blocked in GDB buffer without Triton X-100 for 30 minutes at room temperature. Primary antibodies were diluted in GDB buffer and incubated for 2 h at room temperature. Secondary antibodies and Phalloidin were diluted in GDB buffer and incubated at room temperature for 1 h. The primary antibodies used were rabbit anti-SEPT7 (1:300; IBL America), mouse anti-α-tubulin (DM1α, 1:200; SIGMA). Secondary F(ab’)2 fragment affinity-purified antibodies (1:200) were purchased from Jackson ImmunoResearch Laboratories and included donkey anti-mouse and donkey anti-rabbit antibodies conjugated with AlexaFluor 488 or AlexaFluor 647. Phalloidin conjugated with rhodamine (1:200; Cytoskeleton) or iFluor 647 (1:200; Abcam) were used for actin staining. Samples were mounted in FluorSave mounting medium (EMD Millipore) and imaged with super-resolution structured illumination microscopy using the DeltaVision OMX V4 (GE Healthcare) inverted microscope equipped with an Olympus 60X/1.42 NA objective and sCMOS pco.edge cameras, using immersion oil with a refractive index of 1.515 and a 0.125 µm z-step. Images were reconstructed with the softWoRx software.

**Protein expression and purification**

His-SEPT2/6/7-Strep (SEPT2/6/7) and His-mCherry-SEPT2-SEPT6/7-strep (mCherry-SEPT2/6/7) complexes were expressed as described before (3). *E.coli* BL21 (DE3) bacteria were co-transformed with pnCS-SEPT6/7 (SEPT6/7) and pnEA-vH-SEPT2 or pET15b-SEPT2-mCherry. Bacteria were grown to OD_600_ of 2–3 and induced with 1 mM IPTG (isopropyl β-d-1- thiogalactopyranoside) for 1 h at 37°C for SEPT2/6/7, or to OD_600_ of 0.5 and induced with 0.2 mM IPTG for 16 h at 18°C for mCherry-SEPT2/6/7. Following induction bacterial cultures were centrifuged at ,4000 rpm for 20 minutes at 4°C. Bacterial pellets were resuspended in buffer containing 50 mM Tris-HCl pH 8.0, 150 mM NaCl, 10% glycerol, 10 mM imidazole, 1 mM DTT, 1 mM PMSF, 1 mg/ml lysozyme, 5 µg/ml DNAse I (Millipore) and 1X Bacterial Protease Arrest cocktail (G-Biosciences; 786–330). Bacteria were lysed with sonication, centrifuged at 13,000 rpm for 30 min at 4°C and passed through a 0.45-µm-pore filter. Supernatants were loaded onto gravity flow columns with Ni-NTA agarose beads (Macherey-Nagel; 745400.25) pre-equilibrated with 50 mM Tris-HCl pH 8.0, 150 mM NaCl, 10% glycerol, 10 mM imidazole, 1 mM DTT, 1 mM PMSF. After binding, beads were washed extensively with 50 mM Tris-HCl pH 8.0, 500 mM NaCl, 10% glycerol, 10 mM Imidazole, 1 mM DTT, 1 mM PMSF. Proteins were eluted in buffer containing 50 mM Tris-HCl pH 8.0, 150 mM NaCl, 10% glycerol, 250 mM imidazole, 1 mM DTT, 1 mM PMSF, and were subsequently loaded to a StrepTrap HP column (GE Healthcare) equilibrated with 50 mM Tris-HCl pH 8.0, 150 mM NaCl, 10% glycerol, 10 mM imidazole, 1 mM DTT, 1 mM PMSF. Proteins were eluted in 50 mM Tris-HCl pH 8.0, 150 mM NaCl, 10% glycerol, 10 mM Imidazole, 1 mM DTT, 1 mM PMSF and 2.5 mM d-Desthiobiotin (Sigma; D1411). Septin complexes were dialyzed against 50 mM Tris-HCl pH 8.0, 50 mM NaCl, 10% glycerol, and further purified using an AKTA FPLC system (GE Healthcare) with a Superdex 200 10/300 GL (Amersham Biosciences) gel filtration column.

Rabbit muscle G-actin was labeled with Oregon-green after purification from rabbit muscle acetone powder (Pel-Freeze Biologicals), which was extracted in buffer G (2 mM Tris-HCl pH 8, 200 μM ATP, 0.5 mM DTT, 0.1 mM CaCl2, 1 mM sodium azide). In buffer G extracts, actin was polymerized in the presence of KCl (50 mM) and CaCl_2_ (2 mM), and subsequently depolymerized and purified with size exclusion chromatography (HiLoad 26/600 Superdex 200pg). Oregon-green actin was prepared from polymerized filamentous actin, which was incubated overnight with Oregon Green 488 succinimidyl ester (200 μM; Life Technologies) at 4˚C. Oregon-green-labeled actin filaments were then depolymerized, labeled G-actin was purified by size-exclusion chromatography (Superdex 200 10/300 GL) and stored dialyzed in buffer G.

**Preparation of taxol-stabilized microtubules**

Taxol-stabilized microtubules were prepared by incubating 77% unlabeled tubulin (Cytoskeleton; T240) with 11.5% biotin-tubulin (Cytoskeleton; T333) and 11.5% HiLyte-488-tubulin (Cytoskeleton; TL488M) or 11.5% HiLyte-647-tubulin (Cytoskeleton; TL670M). Tubulin mix (17 µM) was polymerized in BRB80 (80 mM PIPES, pH 6.9, 2 mM MgCl_2_, 1 mM EGTA) supplemented with 10% glycerol, 1 mM GTP and incubated for 30 minutes at 37°C. Then microtubules were supplemented with taxol and were further incubated for 10 minutes at 37°C. After incubation, microtubules were diluted in BRB80 with 10 µM taxol, span for 20 minutes at 100,000×g (Optima TL100; Beckman Coulter) and resuspended in BRB80 with 20 µM taxol.

**In vitro assays of microtubule association with actin filaments**

F-actin (25.5 µM) was made by incubating unlabeled G-actin (Cytoskeleton Inc; AKL99) in the presence of F-actin buffer (20 mM HEPES, pH7.4, 100 mM KCl, 1 mM MgCl_2_, 0.5 mM ATP, and 4 mM DTT) for 1 h at room temperature. F-actin was supplemented with iFluor488 phalloidin (AAT Bioquest; 23115) or rhodhamine phalloidin (Cytoskeleton Inc; PHDR1), and was further incubated for 30 minutes at room temperature. Imaging chambers (3, 10-12) were incubated with 1% Pluronic F-127 for 5 min. Chambers were washed with BRB80, incubated with 5 mg/ml Biotin-BSA (A8549; Sigma-Aldrich) for 5 minutes, followed by a wash with BRB80 and incubation with 0.5 mg/ml Neutravidin (A2666; Invitrogen) for 5 minutes. Subsequently imaging chambers were blocked with BRB80, 1% Pluronic F-127, 1 mg/ml BSA for 5 minutes. Taxol-stabilized microtubules (HiLyte-488-tubulin or HiLyte-647-tubulin) were flowed into the chamber for 15 min. Flow chambers were washed with BRB80, 1mg/ml BSA, 10 µM taxol and incubated with increasing concentrations of recombinant Sept2/6/7 (dark or mCherry-labelled) in BRB80, 1 mg/ml BSA, 10 µM taxol for 15 minutes. After a final wash with BRB80, 1 mg/ml BSA, 10 µM taxol, the reaction mix containing F-actin, 1mg/ml BSA, 10 µM taxol, 0.25% Pluronic F-127, 0.1% κ-casein, oxygen scavenging system (0.5 mg/ml glucose oxidase, 0.1 mg/ml catalase, 4.5 mg/ml d-glucose, 70 mM β-mercaptoethanol) and phalloidin in BRB80 was flowed into the imaging chambers. Imaging chambers were sealed with vacuum grease and still images were taken after 15 min of incubation at room temperature using TIRF microscopy on the DeltaVision OMX V4 imaging platform (GE Healthcare) with 60X/1.49 NA objective (Olympus), sCMSO pco.edge cameras, and the softWoRx software (Fig. 1G-H).

F-actin (25.5 µM) was made by incubating unlabeled human platelet (APHL99; Cytoskeleton, Inc) or rabbit skeletal G-actin (AKL99; Cytoskeleton, Inc) in the presence of F-actin buffer (20 mM HEPES, pH7.4, 100 mM KCl, 1 mM MgCl_2_, 0.5 mM ATP, and 4 mM DTT) for 1 h at room temperature. F-actin was supplemented with X-Rhodamine phalloidin and was further incubated for 30 min at room temperature. F-actin (2 µM) was mixed with taxol-stabilized microtubules (2 µM) (90% unlabeled tubulin and with 10% HiLyte-488-tubulin) in BRB80, 20 µM taxol in the absence or presence of SEPT2/6/7 complexes (500 nM) for 15 minutes at room temperature. Reactions were diluted and an aliquot of each reaction mix was mounted on a slide and sealed with a glass coverslip and nail polish. Samples were imaged on a Zeiss AxioObserver Z1 inverted microscope equipped with a Zeiss a 63x/1.4 NA oil objective, a Hamamatsu Orca-R2 CCD camera and the Slidebook 6.0 software (Fig. 1I).

**In vitro assays of microtubule association with polymerizing actin**

F-actin seeds (5 µM) were prepared by incubating 90% unlabeled rabbit skeletal muscle G-actin with 10% biotin-G actin (Cytoskeleton; AB07) in the presence of KMEI buffer (10 mM Imidazole pH7.0, 50 mM KCl, 1 mM MgCl_2_, 10mM EGTA) supplemented with 1 mM ATP for 1 h at room temperature. F-actin seeds were supplemented with phalloidin iFluor 488 (ATT Bioquest; 23115) or rhodamine phalloidin (Cytoskeleton, Inc; PHDR1) and were further incubated for 30 min at room temperature. After incubation, F-actin seeds span for 20 minutes at 100,000xg. Imaging chambers were incubated with 1% Pluronic F-127 for 5 minutes. Chambers were washed with BRB80, incubated with 5 mg/ml Biotin-BSA (Sigma-Aldrich; A8549) for 5 minutes, followed by a wash with BRB80 and incubation with 0.5 mg/ml neutravidin (Invitrogen; A2666) for 5 minutes. Subsequently imaging chambers were blocked with BRB80, 1% Pluronic F-127, 1 mg/ml BSA for 5 minutes. Taxol-stabilized microtubules (HiLyte-647-tubulin) were flowed into the chamber for 15 minutes. Flow chambers were washed with BRB80, 1mg/ml BSA, 10 µM taxol and incubated with recombinant Sept2/6/7 complexes (dark or mCherry-labelled) in BRB80, 1mg/ml BSA, 10 µM taxol for 15 minutes. Then imaging chambers were washed with KMEI, 1 mg/ml BSA, 10 µM taxol, incubated with F-actin seeds in buffer containing 2 mM Tris pH 8.0, 0.1 mM MgCl_2_, 0.2 mM ATP, 0.5 mM DTT, 1 mg/ml BSA and 10 µM taxol for 10-20 min and finally washed with KMEI, 1 mg/ml BSA, 10 µM taxol. Actin polymerization was initiated by flowing into the chamber actin mix (80% unlabeled G-actin and 20% Oregon green-labeled G-actin) in KMEI supplemented with 0.25% Pluronic F-127, 20 µM taxol, 0.1% κ-casein, 1 mg/ml BSA, 1% methylcellulose and oxygen scavenging system (0.5 mg/ml glucose oxidase, 0.1 mg/ml catalase, 4.5 mg/ml D-glucose, 70 mM β-mercaptoethanol). Imaging chambers were sealed with vacuum grease and imaging was performed every 10 s for 30 minutes at room temperature using TIRF microscopy on the DeltaVision OMX V4 imaging platform (GE Healthcare) with 60X/1.49 NA objective (Olympus), sCMSO pco.edge cameras, the softWoRx software and temperature-controlled chamber (Fig. 2A-D).

**Time-lapse live cell imaging**

B35 neurons (Fig. 2E) were co-transfected with 0.75 µg of pEGFP-Sept7 and F-tractin-miRFP670 using Lipofectamine 2000 (Thermo Fisher Scientific), and 6 h after transfection were switched to differentiation media. Before imaging, cells were washed with Fluorobrite DME medium (ThermoFisher Scientific; A1896701) supplemented with 30 mM HEPES, 0.1 mM dbcAMP, N2 supplement and 1% penicillin/streptomycin/kanamycin (SIGMA and GIBCO). Cells were incubated overnight with SPY555-tubulin (1:10,000 dilution; Cytoskeleton CY-SC203) at 37 ^o^C with 5% CO_2_. Time-lapse image series were corrected for bleaching and denoised using the PureDenoise plugin in Fiji.

Primary rat hippocampal neurons (DIV1; Fig. 2F) were transfected with plasmid encoding for EGFP-Sept7 using Lipofectamine 3000 (Thermo Fisher Scientific). Neurons were imaged 48 h after transfection following an overnight incubation with the Spirochrome dyes SPY555-tubulin and SPY650-FastAct (Cytoskeleton Inc) according to manufacturer's instructions. Images were collected at a rate of 2 s per frame for 2 minutes. Hippocampal neurons were imaged in Gibco Neurobasal medium without phenol red, supplemented with B-27 supplement (Gibco) per manufacturer’s instructions, and 30 mM HEPES.

All time-lapse live cell imaging was performed at 37 ^o^C with the DeltaVision OMX V4 imaging platform (GE Healthcare) using the TIRF module with 60X/1.49 NA objective (Olympus), 488/568/642 nm laser lines, sCMSO pco.edge cameras, the softWoRx software and a temperature-controlled chamber.

**Gene silencing experiments**

Gene silencing experiments were performed in B35 neurons by transfecting 1 μg of plasmid DNAs encoding for mCherry (transfection marker) and scramble control or Sept7 shRNAs with Lipofectamine 2000 (ThermoFisher Scientific). After 24 h, neurons were switched to media for differentiation and after an additional 24 h were processed for immunofluorescence microscopy and SIM imaging (Fig. 2G). Primary rat hippocampal neurons (DIV1) were similarly transfected for 48 h with the GFP-expressing scramble and Sept7 shRNA plasmids using Lipofectamine 2000 as previously described (9). After fixation and staining for microtubules and actin as described above, the growth cones of shRNA-expressing (GFP-positive) neurons were imaged with an inverted Zeiss AxioObserver microscope with a Zeiss oil 63X/1.4 NA objective, a Hamamatsu Orca R2 CCD camera and the Slidebook 6 software (3i; Intelligent Imaging Innovations).

**Image and statistical analyses**

Quantification of fluorescence by line scans (Fig. 1B) was performed in Fiji using the plot profile function, and fluorescence intensities were plotted as fraction of the maximum intensity, which was assigned to a value of 1. Quantification of the fluorescence intensity of phalloidin-stabilized actin filaments along microtubules (Fig. 1G-H) was performed in Fiji by manually drawing 5 pixel-wide line along the length of taxol-stabilized microtubules, and deriving the average value of actin fluorescence after subtraction of the background fluorescence using the subtract background tool in Fiji (rolling ball radius, 30 pixels). Overlap between growing actin filaments and taxol-stabilized microtubules (Fig. 2B) was quantified after 30 minutes of actin polymerization in the absence or presence of Sept2/6/7. Using fluorescence intensity thresholding, individual actin filament and microtubule masks were made after subtraction of background fluorescence (rolling ball radius, 50 pixels). Using the AND function of Fiji, overlapping areas between actin and microtubule masks were derived, and their surface areas were calculated as percentage of the total area of actin filaments. The percentage area of total actin overlapping with microtubules was calculated as the fold change between the absence and presence of Sept2/6/7 with 1 being the overlap in the absence of Sept2/6/7 (Fig. 2B). The collision angle of each polymerizing actin end with a microtubule lattices was measured in Fiji using the angle tool, and their distribution was plotted as percentage of total collision events in increments of 5 degrees between 0^o^ to 90^o^. The outcome of each collision was categorized by visual inspection for coalignment with the microtubule lattice (overlap) or non-overlapping crossover, and plotted as percentage of total collisions in their respective 5^o^ angle increments (Fig. 2C).

Quantification of actin growth rates was performed in Fiji. Time-lapse image series were corrected for bleaching using the bleach correction plug-in. Kymographs were generated using the KymographBuilder plug-in. Growth rates were derived by manual segmentation of the trajectories of growing actin filaments. The velocity measurement tool macro was used to calculate growth rates. Microtubules with multiple and simultaneous actin polymerization events were excluded from the analysis.

Growth cones were categorized by visual inspection into fan-shaped, club-shaped or collapsed, and each category was expressed as percentage of total growth cones (Fig. 2G). Growth cone surface areas were quantified in Fiji after boosting the intensity levels of the phalloidin channel, which enabled delineation of the membrane edges and tracing of the surface area.

All statistical analyses were performed in GraphPad Prism software. Data sets were tested for normal distribution using the Kolmogorov–Smirnov test. Mean, SEM, and p values were derived using Student’s t-test for normally distributed data and the Mann–Whitney U test for non-normally distributed data. Analysis of variance (ANOVA) was used for multiple comparison of groups. For normally distributed data, ordinary one-way ANOVA was used followed by the Tukey’s multiple comparison test. For non-normally distributed data, the Kruskal–Wallis H test was used followed by the Dunn’s multiple comparison test.

**Movie S1**

Polymerizing actin filaments cross over microtubules in the absence of septins. Representative movie of TIRF imaging of actin (green) polymerizing from phalloidin-stabilized actin seeds (magenta) in the presence of taxol-stabilized microtubules (red). Images were acquired at the rate of 5 s per frame. Scale bar, 5 μm. Also see Figure 2A-B.

**Movie S2**

Capture and growth of polymerizing actin filaments on the lattice of microtubules in the presence of Sept2/6/7. Representative movie of TIRF imaging of actin (green) polymerizing from phalloidin-stabilized actin seeds (magenta) in the presence of taxol-stabilized microtubules (red), which were pre-coated with recombinant Sept2/6/7 complexes (dark; 500 nM). Arrowheads point to microtubule sites of actin filament end docking and overtaking, marking the beginning and end of actin-microtubule zippering. Images were acquired at the rate of 5 s per frame. Scale bar, 5 μm. Also see Figure 2D.

**Movie S3**

Actin filament growth on the lattice of a microtubule coated with mCherry-Sept2/6/7. Representative movie of TIRF imaging of actin (green) polymerizing from phalloidin-stabilized actin seeds (not shown) in the presence of taxol-stabilized microtubules (red), which were pre-coated with recombinant mCherry-Sept2/6/7 complexes (cyan; 500 nM). Arrowheads point to the microtubule sites, on which actin zippering along the microtubule lattice begins and ends. Images were acquired at the rate of 5 s per frame. Scale bar, 5 μm.

**Movie S4**

Actin filament growth at a Sept7-enriched site of growth cone microtubules. Time-lapse TIRF imaging of an elongating actin filament (cyan; F-tractin-miRFP670) that originates from a microtubule (orange; SPY555-tubulin) associated Sept7 (magenta; GFP-Sept7) in the growth cone of a differentiated B35 neuron. The movie begins at a viewing rate of 10 frames per second and slows down to 2 frames per second for visualizing the stages of actin filament growth (arrowhead). Scale bar, 2 μm. Also see Figure 2E.
